# A Novel Rat Spatial Inference Task Reveals Rapid Deduction of Transitive Relationships Using Schemas and Deliberation

**DOI:** 10.1101/2025.08.28.672785

**Authors:** Blake S. Porter, Catherine Shi, Evgeniia Kozlova, Shantanu P. Jadhav

## Abstract

Inferential reasoning is a vital cognitive ability that enables animals to navigate novel situations by leveraging existing relational knowledge of memory schema, with hypothesized roles of prefrontal cortical – hippocampal circuits. Transitive inference (TI) tasks test the ability of subjects to infer relationships within a value hierarchy (e.g., A>B>C>D>E) after being trained only on adjacent premise pairs (e.g., A-B, B-C, etc.). In rodents, current TI paradigms are primarily based on odor-cues and have several limitations that preclude investigation of physiological mechanisms underlying schemas and deliberation. To address these challenges, we developed a novel, automated spatial TI task for rats using a radial maze with maze arms as premise elements and a dedicated deliberation zone. Most rats successfully learned the premise pairs over training. Further, animals demonstrated rapid, successful inference (test pair B>D and control pair A>E) within a single test session, with higher initial accuracy than comparable premise pairs, indicating the use of schema-based inference. We also investigated vicarious trial and error’s (VTE), a behavioral correlate of spatial deliberation. VTE behavior was elevated on choice trajectories early in learning, when novel premise pairs were introduced, and generally for incorrect trials, corresponding to the hypothesized association of VTE’s with uncertainty. Further, rats also exhibited elevated VTE behavior with high variability during inference testing, with individual variability suggestive of varying strengths of schema usage. Our findings demonstrate the feasibility of a rodent spatial TI task that provides new insights into the behavioral correlates of schemas and deliberation for inferential reasoning.

---

In an ever-changing world, animals will encounter novel situations where they must rely on existing knowledge to make the best decision. Inferential reasoning is the cognitive ability to draw upon existing knowledge that shares common features with a novel scenario to make informed decisions. Memory schemas, or relational networks of abstract and generalized information, likely aid inferential reasoning by enabling the retrieval of information that matches elements of a novel situation (Varga et al., 2024). Leveraging the relational network of schemas enables animals to infer potential action plans and their outcomes without having to explicitly experience them first (Tenenbaum et al., 2011). Thus, inferential reasoning not only saves time and energy, but it can reduce risk by avoiding inferred harm.

Transitive inference (TI) is a hallmark inference paradigm that tests a subject’s ability to infer the relationships of elements in a value hierarchy (e.g., A>B>C>D>E; Vasconcelos, 2008). Commonly, subjects are only trained on adjacent items of the set (e.g., A-B, B-C, etc.; also known as premise pairs), with inference testing consisting of novel combinations (e.g., B-D). Thus, to successfully infer, subjects must form a memory schema of the complete value hierarchy built up from the premise pairs.

Transitive inference appears to be a highly conserved cognitive function as it has been demonstrated in insects (Tibbetts et al., 2019), fish (Grosenick et al., 2007), rodents (Davis, 1992; Dusek & Eichenbaum, 1997), multiple bird species (Lazareva et al., 2004; Steirn et al., 1995; von Fersen et al., 1991), non-human primates (NHP; Brunamonti et al., 2016; Gillan, 1981; Mcgonigle & Chalmers, 1977; for review: Vasconcelos, 2008), and humans (Bryant & Trabasso, 1971; Preston et al., 2004).

Transitivity is an important property of relationships between stimuli, actions, and information more generally. Utilizing inferential reasoning on transitivity allows animals to understand the world around them without having to experience every possible relationship (Eichenbaum, 1999, 2000; Tenenbaum et al., 2011). An ethological example of the benefits of transitive inference is social hierarchies (Becker et al., 2021). By observing peers fighting and considering their own rank, territorial beta fish utilize transitive inference to determine their social rank and abilities to fight other peers (Grosenick et al., 2007). Being able to infer their rank can help them avoid obvious defeats and take on easier rivals.

However, we know little about the underlying physiological mechanisms that support transitive inference. Computational modeling using artificial neural networks has demonstrated that transitive inference can occur in many model architectures (Frank et al., 2003; Kay et al., 2024; Kumaran & McClelland, 2012; Miconi & Kay, 2025; Wu & B Levy, 2001; Wu & Levy, 1998), including hippocampal network models (Frank et al., 2003; Kumaran & McClelland, 2012; Wu & B Levy, 2001; Wu & Levy, 1998). Human fMRI studies (Acuna et al., 2002; Heckers et al., 2004; Schlichting & Preston, 2016; Zalesak & Heckers, 2009; Zeithamova et al., 2012) and human (Koscik & Tranel, 2012; Wing et al., 2021) and animal lesion studies (DeVito, Kanter, et al., 2010; DeVito, Lykken, et al., 2010; Dusek & Eichenbaum, 1997) have demonstrated the necessity of the hippocampus and prefrontal cortices in transitive inference. Recent electrophysiological studies with non-human primates have provided some key insights into how prefrontal cortical circuits aid in inferential reasoning (Brunamonti et al., 2016; Ramawat et al., 2022, 2023). Sleep is also known to play an important role in consolidation of schema-based memories (Abdou et al., 2024; Durrant et al., 2015; Ellenbogen et al., 2007; Santamaria et al., 2024). Schemas and inference are thus known to involve key roles of prefrontal cortical and hippocampal circuits in the mammalian brain. Rodent models offer the possibility of electrophysiological investigation of neural ensemble mechanisms across distributed circuits underlying schema formation over the course of waking and sleep states during learning, as well as mechanisms underlying deliberation and deduction during inference testing. However, studying these neurophysiological processes has been challenging in rodent models due to several disadvantages of commonly used TI paradigms.

Most rodent inference studies have relied on odor-cue based paradigms requiring rodents to discriminate between pots filled with scented sand where each odor is an item in the value hierarchy (e.g., A is paprika, B is coffee, etc.; André et al., 2012; Davis, 1992; DeVito, Kanter, et al., 2010; DeVito, Lykken, et al., 2010; Dusek & Eichenbaum, 1997; Van der Jeugd et al., 2009; Van Elzakker et al., 2003). This setup makes it challenging to pinpoint when exactly rodents are deliberating and potentially utilizing inferential reasoning. Furthermore, many of the previous TI paradigms required the experimenter to manually set up each trial, which slows down throughput and has obvious disadvantages for physiological investigation. Other memory schema tasks, such as spatial odor-cue based paradigms have been described, but remain unreplicated and physiologically intractable (Tse et al., 2007, 2011; Wang et al., 2012). Encoding of odor, spatial-context, and value has been investigated in hippocampal and orbitofrontal cortex (OFC) schemas (Farovik et al., 2015; McKenzie et al., 2014, 2016), and OFC schemas have been described in odor sequence tasks (Zhou et al., 2020; Zong et al., 2025), however, odor-based schema representations in prefrontal and hippocampal circuits are still relatively under-studied compared to spatial representations. In particular, the nature of coherent, global schema representations based on prefrontal-hippocampal interactions, posited to be a central mechanism for memory schema and inference (Varga et al., 2024), require simultaneous recordings in both regions during behavior and sleep across learning (Lin & Zhou, 2024; Shin & Jadhav, 2024; Tang et al., 2023). Further, investigation of the potential role of sleep mechanisms in consolidation and schemas (Abdou et al., 2024; Durrant et al., 2015; Ellenbogen et al., 2007; Santamaria et al., 2024), such as sharp-wave ripple replay events (Barron et al., 2020; for review: Tang & Jadhav, 2019), can benefit from related studies in spatial memory tasks

To address these issues, we sought to develop a novel, automated spatial transitive inference (TI) task for rodents to leverage rats’ natural abilities to forage for food and to be amenable to physiological investigation. Using maze arms on a radial maze as hierarchy elements, rats can be trained on premise pair relationships (A>B, B>C, C>D, D>E) over multiple training sessions with interleaved behavior and sleep states that are amenable to investigation of mechanisms underlying schema formation over the course of learning. Premise pair learning is then followed by tests of inferential reasoning along with control tests, with successful inference highlighting the use of schemas and deliberation, amenable to investigation of mechanisms underlying inference based on prior knowledge encoded in schemas.

We also designed the paradigm with a delineated area of the maze where rats can deliberate choices before execution, facilitating the elucidation of the behavioral and neural correlates of inferential reasoning. A promising approach to quantify deliberative behaviors is vicarious trial and error (VTE; for review: Redish, 2016). Vicarious trial and error events were first described by Gentry and Muenzinger (Muenzinger, 1956) and popularized by Tolman (Tolman, 1939, 1948) as a potential behavioral readout of deliberation where rats vicariously test specific action plans and their outcomes before making a choice. This theory has been bolstered by evidence from neural recordings showing hippocampal-prefrontal oscillatory interactions (Amemiya & Redish, 2018; Rosenblum et al., 2025; Stout et al., 2022) and hippocampal theta sequences sweeping ahead of animals to multiple potential routes during VTEs (Johnson & Redish, 2007; Papale et al., 2016). Yet the exact role of VTEs and their prevalence remains unclear.

VTEs can be most prevalent during learning while rats deliberate during periods of uncertainty, and then decline as a task becomes habitual (Gardner et al., 2013; Packard & McGaugh, 1996; Regier et al., 2015; Smith & Graybiel, 2013). However, VTEs may also persist in especially challenging tasks even as the rats become proficient, possibly reflecting the ongoing need for deliberation (Miles et al., 2024; Steiner & Redish, 2014; van der Meer et al., 2010). Thus, our development of a spatial transitive inference task with a deliberation zone allows investigation of the role of VTEs in uncertainty and deliberation during both initial learning of premise pairs and during later tests of inferential reasoning, providing a foundation for neurophysiological investigation of the mechanisms underlying deliberation and inferential reasoning.

## Methods

### Subjects

All experimental methods were approved by the Brandeis University Institutional Care and Use Committee (#21001 and #24001-A) and conducted under the guidelines of the US National Institutes of Health. A total of 14 adult male Long-Evans rats (weighing between 350 – 536 grams) were used. Five of the rats were from a TH-Cre line (Witten et al., 2011) and used in a previous experiment (Ding et al., 2025) before the transitive inference task. All other rats were obtained from Charles River Laboratories or bred in-house.

Initially, animals were dual housed and given ad libitum access to food and water on a 12-hour/12-hour light/dark cycle (7am-7pm light), followed by single housing for food restriction. Before training began, rats were food-restricted to no less than 85% of their free feed weight. The rats were habituated to the experimenters and evaporated milk (Nestlé S.A.) before being trained to run back and forth on a linear track.

### Linear Track Pre-training

Rats were pre-trained to run on a linear track. The linear track was placed in the same room as the main experimental radial maze and constructed of the same maze components. An 8-arm radial maze, with automated rule implementation, reward delivery, and barrier control (Porter et al., 2025) was used for the TI experiment. While running on the linear track, pneumatic barriers were randomly triggered on the radial maze to habituate the rats to the sound. A lick requirement to dispense the reward (evaporated milk) was introduced on the linear track, with rats trained to lick three times before a reward was automatically dispensed. This lick requirement ensured that 1) there was a deliberate behavioral readout of animals’ choice beyond just traveling to a reward well, 2) the reward was not pre-dispensed so rats could not navigate by smell or the sound of the pump to the correct radial-maze arm, and 3) it avoided rats accidentally dispensing a reward (e.g., by stepping on the lick sensor). Once readily shuttling on the linear track (∼ 5 rewards/min), non-implanted animals and the already implanted TH-Cre animals moved onto premise pair training. One more rat (Long Evans) was implanted with a 64-tetrode microdrive targeting the prefrontal cortex and the hippocampus as described in Ding et al. (2025).

### Premise Pair Training

Each maze arm used as a premise element had its own unique visual and textural cues such as black foam, hot glue dots, or vinyl stripes (**Supplement Figure 1A**). Colorful shapes were adhered to the walls of the experiment room for distal visual cues. Arms were separated by 14” high walls spanning 24” from the center so that rats could not jump from one arm to another. Arm access was controlled by 12” pneumatic barriers situated within the track joints between the maze center and each radial arm. At the end of each arm was a reward well that would dispense an evaporated milk reward (∼ 0.1mL) after rats licked three times.

One arm was designated as the Home arm and this arm was kept constant across all rats. For each rat, five arms were randomly assigned to the five items of the value hierarchy (i.e., A, B, C, D, E). Any arm assignments that were assessed to have a simple egocentric strategy that could be used to successfully solve the task (e.g., always go to the left- or right-most arm) were not used and new assignments were generated until satisfactory. A rat’s arm assignments stayed the same throughout all of training and testing (assignments for all animals shown in **Supplement Figure 1B**).

The transitive inference trial structure was as follows (**Figure 1D**): rats always began each trial at the Home arm reward well where they received a reward. Upon licking at Home, the barriers were automatically set for the next trial to allow access to the two relevant arms for the trial’s item-pair.

**Figure 1:**
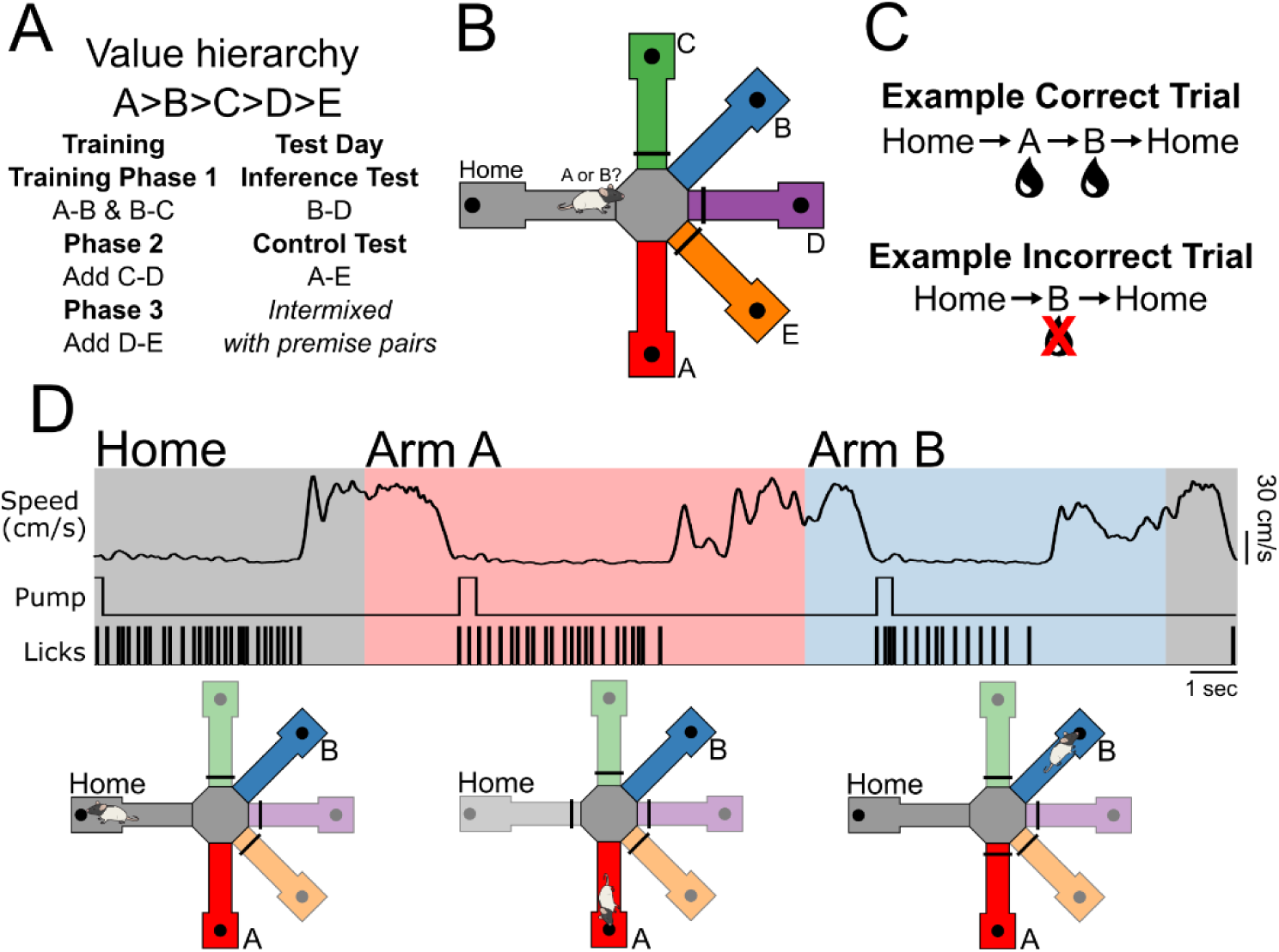
Spatial transitive inference paradigm. **A)** Paradigm overview of training and testing schedules. **B)** Maze schematic depicting an example A-B trial where rats exit home and must decide which is the correct arm to visit. Other arms/items are blocked off by barriers (black bars). Black circles represent reward wells. **C)** Examples of correct and incorrect trial structure for an A-B trial. **D)** Top: Timeline of one example A-B trial showing speed, reward Pump, and licks at reward wells. Animals start at Home to initiate a new trial. Rats then travel to the center and decide on which arm to choose. In this trial, the rat is correct and consumes a reward at arm A reward well before traveling to arm B for a second reward, and then returning Home. **Bottom:** schematic showing the state of the barriers and arm availability over the course of the trial. Greyed-out arms are blocked by barriers (black bars).

**(i) Correct trials**: Rats would enter the center of the radial maze and, given the two open arms, needed to choose the correct one based on the latent value hierarchy (A>B>C>D>E). If rats chose correctly (e.g., arm A when presented with A-B), they would be rewarded at the reward well at the end of the correct arm (i.e., A), and the Home arm would be automatically blocked off with a barrier. Rats would then have a free choice with only the second arm (i.e., B) available as an open arm, for a second “free” reward. Once the sequence of A followed by B visits were complete, this lowered the Home barrier and raised the first arm barrier (e.g., A), so that only return to Home arm was available. Thus, for correct trials, rats receive rewards on both arms comprising the premise pair, with the critical choice being the first arm of the pair. We chose to reward the second arm visited in the premise pair for correct trials to maintain motivation and to keep reward conditions the same regardless of whether an arm would be the first or second in a premise pair (e.g., C in B-C and C-D), unique from previous versions of rodent TI (André et al., 2012; Davis, 1992; DeVito, Kanter, et al., 2010; DeVito, Lykken, et al., 2010; Dusek & Eichenbaum, 1997; Van der Jeugd et al., 2009; Van Elzakker et al., 2003).

**(ii) Incorrect trials**: If the animal chose incorrectly at the beginning of the premise-pair trial (e.g., the animal chose arm B when presented with A-B), rats would receive no reward at the incorrect arm (i.e., B) and the barrier to the correct arm (i.e., A) would be raised so it could not be visited. Thus, for incorrect trials, there was no reward and rats had to return Home to initiate the next trial.

Rats received no pre-exposure or pre-training to the 8-arm maze. Rats were trained once per day for five to seven days per week. Training sessions lasted at least fifteen minutes but were stopped at 60 minutes. After fifteen minutes, the session could also be ended if the rat did not attempt a trial for two minutes. All rats developed a habit of sitting and waiting at the Home arm to end the session.

The premise pair training proceeded in three phases (**Figure 1A**), and rats progressed to the next phase when they met the criterion of performing 75% or more correct for all premise pairs in the current phase for two days in a row. This criterion was chosen to be similar to previous rodent TI tasks (DeVito, Kanter, et al., 2010; Dusek & Eichenbaum, 1997; Silverman et al., 2015; Van der Jeugd et al., 2009; Van Elzakker et al., 2003). (i) For the first phase of training, rats were trained on the premise pairs A-B and B-C. On each trial, a premise pair was randomly chosen (A-B or B-C), and animals had access to only the two arms of the current premise pair. (ii) The second phase added C-D, with each trial randomly chosen to be either A-B, B-C, or C-D, and (iii) the third phase added D-E, and thus had all four premise pair trial types (A-B, B-C, C-D, D-E trials).

By default, premise pairs were randomly intermixed. However, to avoid random long streaks and to facilitate learning, if one premise pair had three fewer rewards than any of the other premise pairs, that lower-reward premise pair was presented until its reward count caught up to the others. Thus, if rats were struggling to learn certain premise pairs, those pairs would be presented more often. On occasion, if a rat was severely struggling with certain premise pairs, a training day would begin with fixed blocks (∼ 5-10 trials long) of only those premise pairs before the intermixing of all premise pairs.

These block sessions were counted in the total training days but were never used for determining the training criterion; only fully intermixed sessions were used to determine if a rat was ready for the next training phase or for testing. Once a rat reached the final criterion of performing 75% or better on all four premise pairs for two days in a row, inference testing commenced the following day.

### Transitive Inference Test Day

Inference testing was conducted for just one day. While rats may be capable of learning the new pairs within a single day, this is unlikely given the training times of premise pairs (**Figure 2**); however, repeated exposure to the test pairs over multiple days can result in them learning them through trial and error rather than inferring their choices. The single test day session allowed us to compare initial performance on the test pair B-D with initial performance on the premise pairs during training. On test day, rats were presented with a total of six item-pairs, the four premise pairs (A-B, B-C, C-D, and D-E), the inference test pair (B-D), and a novelty, end-item control pair (A-E). The reinforcement schedule was the same as training for all pairs, i.e., correct B-D and A-E choices were rewarded along with all premise pairs (Vasconcelos, 2008). Pair presentation was controlled by randomly generating blocks of six trials where each item-pair was presented once in the block. This ensured a roughly equal number of trials across pairs. The first two blocks were only the four premise pairs; all subsequent blocks were six pairs.

**Figure 2:**
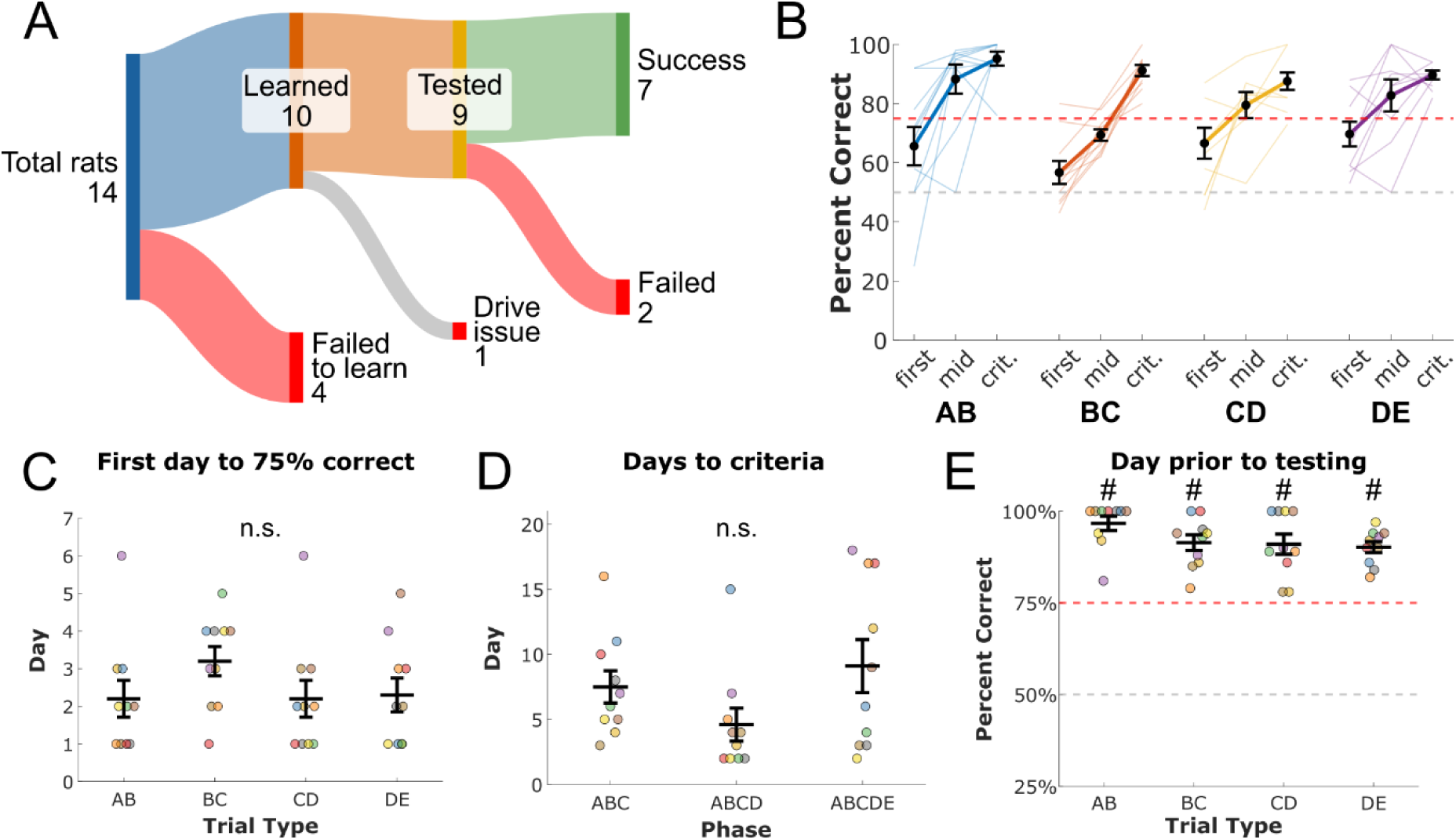
Transitive inference training. **A)** Sankey diagram of rat training and testing progression. **B)** Performance over training for premise pairs. Performance on the first training day, the final criteria day for a training phase, and a midpoint day are shown (n=10 rats). **C)** Number of days to first achieve 75% correct performance in a behavioral session for the given trial type. **D)** Number of days to finish a training phase by reaching 75% correct for each trial type for two days in a row. **E)** Rats’ performance on all premise pairs on the training day before the test day. **C-D**. Each color represents data from one rat, colors consistent across all figures. n.s., p > 0.05 for one-way ANOVA across trial types. # indicates premise pair performance is significantly greater than 75% correct (one-tailed, one-sample t-test, n=10), with Bonferroni corrected p value (p < 0.0125).

### Behavioral Analysis

All data was collected with SpikeGadgets’ environmental control unit (ECU) and their StateScript software. An overhead camera (Mako G-158C, Allied Vision Inc.) synced to SpikeGadgets recorded video at 30 frames per second. Rats’ position was tracked with DeepLabCut (DLC; Nath et al., 2019). For non-implanted animals, the nose was tracked for position, and for implanted animals, the light-emitting diodes on their implant were tracked for position. To ensure accurate tracking, any DLC tracked position with a likelihood less than 0.98 was removed and a linear interpolation was applied. A boxcar filter of 5 frames (∼ 0.17 seconds) was then applied to remove jitter.

### Training Day Selection

Rats had a highly variable number of training days (shown in **Figure 2C-D**). In order to compare performance (**Figure 2B**) and VTEs (**Figure 4D & E**) across training, we divided the premise pair training times into thirds. The first day was selected as the first day a premise pair was presented. The criterion day was the day before rats move onto the next training phase (for A-B, B-C, and C-D) or the day before test day (for D-E). The midpoint day was determined by the day between the first and the criterion day, rounded up to a day if needed. Some rats did not have a midpoint day due to meeting the criterion in just two days. In these rare cases, the midpoint day was represented as a “not a number” (NaN). For all days, if a day had fewer than five trials, it was not used, and the day search was iterated until a day with more than five trials was found. This was primarily done to avoid the earliest training days for some rats who were slow to habituate to the maze and attempted very few trials.

### Vicarious Trial and Error and zIdPhi metric

In order to quantify VTEs, we used the standard integrated absolute change in angular velocity of the head (Idphi) metric developed by Papale et al. (2012), which was applied to choice trajectories from Home to the first arm of choice. For every recording session, a region of interest was manually drawn for the center area of the maze as well as for every arm. The center zone comprised the area inside all the arm barriers. The arm zones were drawn at the barrier to the length of the arm. A choice trajectory was determined from the frame that the rat (nose or LED) entered the center zone to the frame that they entered an arm that was not Home. If rats entered the center and then re-entered the Home arm, the trial was not analyzed. If rats entered a choice arm but did not lick at the reward well and went to the other choice arm, the trial was also not analyzed. Trajectories longer than two seconds in the center zone were also excluded so that behaviors such as grooming or excessive exploration in the center were not analyzed.

Choice arms were all at different angles from the Home arm. Two arms were at ±90°, two arms were at ±135°, and one arm was 180° from Home. To account for this, Idphi values were z-scored separately for each arm and for each animal. This ensured the relatively low baseline Idphi values for the 180° arm and relatively high values for the 90° arms did not bias the mean or standard deviation for other arms. Z-scoring was done independently for each rat using all their trajectories from both training and test to determine zIdPhi values for analysis.

### Statistical Testing

Parametric tests were used unless otherwise stated. Error bars show the standard error of the mean (SEM) unless otherwise stated. Significance was tested with an alpha of 0.05. Bootstrapping for testing correct versus incorrect trials was run 100 times using MATLAB’s datasample function with replacement.

### Transparency and Openness

Materials, data, and analysis code for this study are available by emailing the corresponding author. Data were analyzed using custom MATLAB (R2021b) code including code provided by the Redish lab for the IdPhi analysis (personal correspondence). Animals’ position was determined using DeepLabCut (Nath et al., 2019). This study’s design and its analyses were not preregistered.

## Results

### Transitive inference paradigm

We used a five-item transitive inference set (A>B>C>D>E; **Figure 1A**) where arms of an automated eight-arm radial maze (Adapt-A-Maze, Porter et al., 2025) represented premise elements/items of the set (**Figure 1B**). The task was completely automated, with arm availability based on trial type and controlled by pneumatic barriers (**Figure 1D**). Specifically, for a given trial type, only the arms associated with the items of the current pair were kept open by lowering the barriers (e.g., A-B, only the arms for A and B were available with all other arms’ entries closed with opaque barriers). One arm served as the Home arm with a Home reward well where rats needed to return after every trial (correct or incorrect) to initiate the next trial. For each rat, item identity was randomly assigned to arms (configurations for each animal shown in **Supplemental Figure 1B**). However, configurations were rejected if they were solvable by an obvious egocentric strategy (e.g., “always go left”) and re-generated until satisfactory. A given animal’s item-arm configuration was constant for the entire experiment.

Rats were trained on the inference set iteratively by adding new premise pairs once performing at criterion on the existing pairs, with each trial corresponding to a randomly chosen premise pair, and pairs presented in an intermixed manner across trials (**Figure 1A**). Training began with intermixed presentations of A-B and B-C trial types. We used a criterion of 75% correct for all present pairs for two days in a row to be considered proficient. Once the criterion was reached, the next premise pair was added to the existing premise pairs until all four premise pairs were present (i.e., A-B, B-C, C-D, and D-E). This design allows us to ask questions about schema accommodation and assimilation as new pairs are added, since the existing task schema likely needs to be updated to accommodate new pairs.

For correct trials, rats visited the second item in the premise pairs as a free rewarded choice after the visiting the first item, e.g., a correct A-B trial consisted of rats getting a reward for correctly choosing A when presented with open arms A and B, then another reward at B (**Figure 1C & D**). Thus, animals were rewarded at both items in the hierarchy in the correct order. For incorrect trials, if rats made the wrong choice (e.g., B for an A-B trial), no reward was received, and animals needed to return Home to start a new trial (i.e., arm A would be blocked off once B was chosen). We chose this reward schedule in order to avoid the end item (i.e., E) never being rewarded, and potentially never being visited during periods of high performance. This task design is distinct from previous odor-based rodent TI tasks (André et al., 2012; Davis, 1992; DeVito, Kanter, et al., 2010; DeVito, Lykken, et al., 2010; Dusek & Eichenbaum, 1997; Van der Jeugd et al., 2009; Van Elzakker et al., 2003), which introduced all premise pairs in blocks of trials and with no reward for choosing the second item in a premise pair for correct trials; this design was for our aim of investigating memory schema assimilation during learning, and to avoid potential confounds of no rewards or visits for end item of a set.

### Premise Pair Learning

A total of 14 rats were trained on the transitive inference paradigm (**Figure 2A**).

Of these 14, four failed to learn the premise pairs. Failure to learn was not operationalized, but was characterized by rats reaching more than 30 days of training with no consistent progress towards reaching the final criterion for all four premise pairs. These rats would often have stochastic day-to-day performance, varying from performing greater than 75% for a pair on one day to below chance on the same pair the next day. All four of these rats did finish phase 1 of training (A-B, B-C) and thus understood the general task structure. However, adding the subsequent pairs seemingly proved too challenging. The data from the four rats who failed to learn were not analyzed.

Ten rats successfully learned all four premise pairs and performed at least 75% correct on each of the four pairs for two days in a row. Due to all rats having a different number of training days, we divided the days each premise pair into an early day it was presented (“first”), the criterion day (“crit.”) before the next phase (next training phase or test day), and a midpoint day (“mid”) between the two (**Figure 2B**). Considerable variability was seen across different rats on time to learn specific premise pairs, though no one pair was significantly different from the others (F(3,36) = 1.13, p = 0.35; **Figure 2C**). Most rats obtained 75% correct on A-B within the first 1-2 days (mean 2.2 ± 1.6 days) as they essentially just needed to learn to go to A for this end-item pair. In contrast, B-C tended to have a longer time (3.2 ± 1.2 days) to achieve 75% for the first, time as rats needed to overcome their avoidance of B from A-B in order to get the premise pair correct. Afterwards, rats picked up C-D (2.2 ± 1.6 days) and D-E (2.3 ± 1.4 days) relatively quickly. However, when a new premise pair was added, the performance on other premise pairs tended to temporarily fall, possibly reflecting schema accommodation and the need to update the previous end item to now be a correct choice in the right context (e.g., C for B-C when C-D is added).

There was no significant difference in the average total training days for each training phase (F(2,27) = 2.1, p = 0.14; **Figure 2D**). However, a great deal of variance can be seen across rats, especially for the final phase with the full set (A-B, B-C, C-D, D-E). Some rats seemed to be able to quickly integrate D-E within a few days, while others needed upwards of 18 days to reach the criterion of 75% correct for all four pairs for 2 days in a row. On average, rats took 21.2 days (standard deviation (S.D.) = 10.6) to complete all training, consisting of an average of 2,215 total trials (S.D. = 737).

Despite the variability in training times, all rats who learned the task performed ∼90% correct on all four premise pairs on the final training day with no significant difference across pair performances (F(3,36) = 1.89, p = 0.15; **Figure 2E**).

### Transitive Inference Test Performance

One animal who successfully learned the task had a drive implant issue and could not be tested. The remaining animals (N = 9) were tested on their ability to infer a novel pair of items, B-D. We also tested rats’ ability to perform on A-E, which serves as a novelty control and does not require inference to solve as it consists of both end-items (Davis, 1992; Dusek & Eichenbaum, 1997). Animals were thus presented with six pairs in a random order on test day, four from the training set (A-B, B-C, C-D, D-E), one test pair B-D, and one control pair A-E, with at least 10 trials for each pair. Overall, there was no significant difference in performance across these trial types on test day (F(5,48) = 0.83, p = 0.54; **Figure 3A**). Eight out of the nine rats tested inferred B-D, as shown by their average performance on B-D being around 85% correct. The rat who failed the inference test did worse than chance (∼33% correct), possibly indicating a strong bias to arm D when faced with uncertainty. A second animal did infer B-D, however, failed the A-E control, as well as at A-B trials, performing at chance levels. This suggests that the animal did not possess a schema of the value hierarchy. We consider both animals as failing the TI test since either test or control pair performance was below criterion level. Further performance analyses were performed on N=7 animals that achieved criterion performance on both the inference test and control pair.

**Figure 3:**
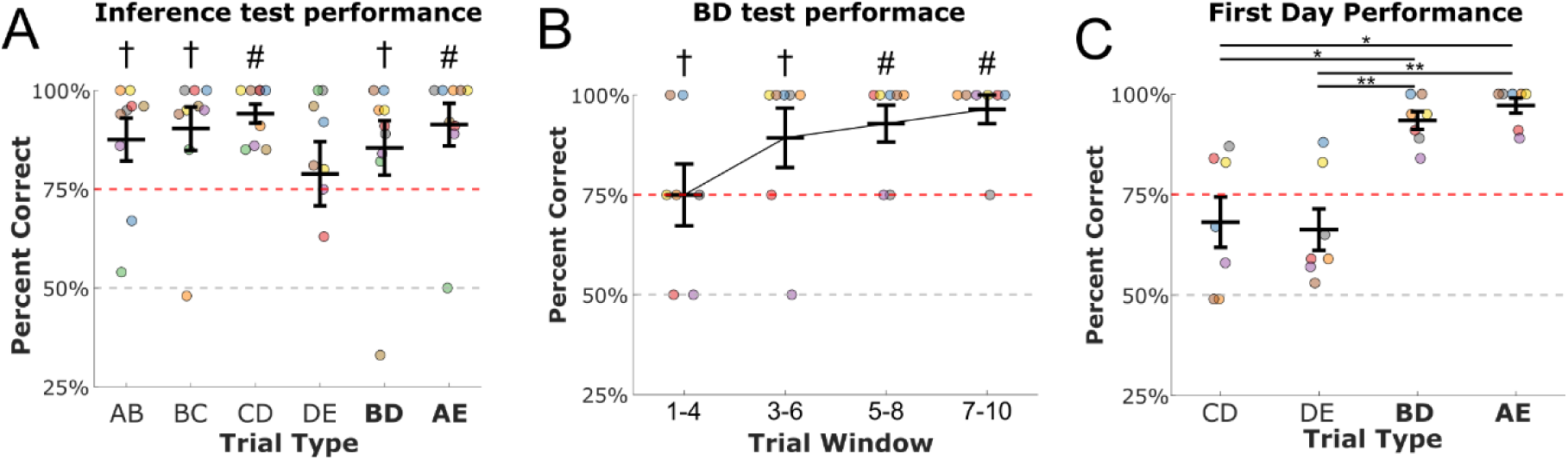
Inference testing. **A)** Performance on testing day for n=9 rats. As a group, rats performed well above chance level on all trial types with no significant difference across trial types. However, one of the eight rats failed to infer the test pair B-D and one failed to infer the control pair with end-items A-E. Four rats showed <75% criterion performance on original premise pairs on test day, despite >75% criterion performance on previous day. **B)** Performance of rats who successfully inferred B-D and A-E, shown for the first 10 B-D trials (n=7 rats). A 4-trial window was used with a 2-trial overlap. **C)** First-day performance of rats who successfully inferred, for premise pairs C-D and D-E on the first day of presentation with the trial type. Note that B-D and A-E is same as seen in **A** for the subset of n=7 rats. Rats showed significantly performance on B-D and A-E test pairs than on the original premise pairs on first day of that trial type. Each color represents data from one rat, colors are consistent across all figures. Error bars show S.E.M. † indicates significantly greater than 50%, # indicates significantly greater than 75%, one-sample t-test with Bonferroni correction. *, p < 0.05; **; p < 0.01 for post-hoc pairwise comparisons from one-way repeated measures ANOVA.

**Figure 4:**
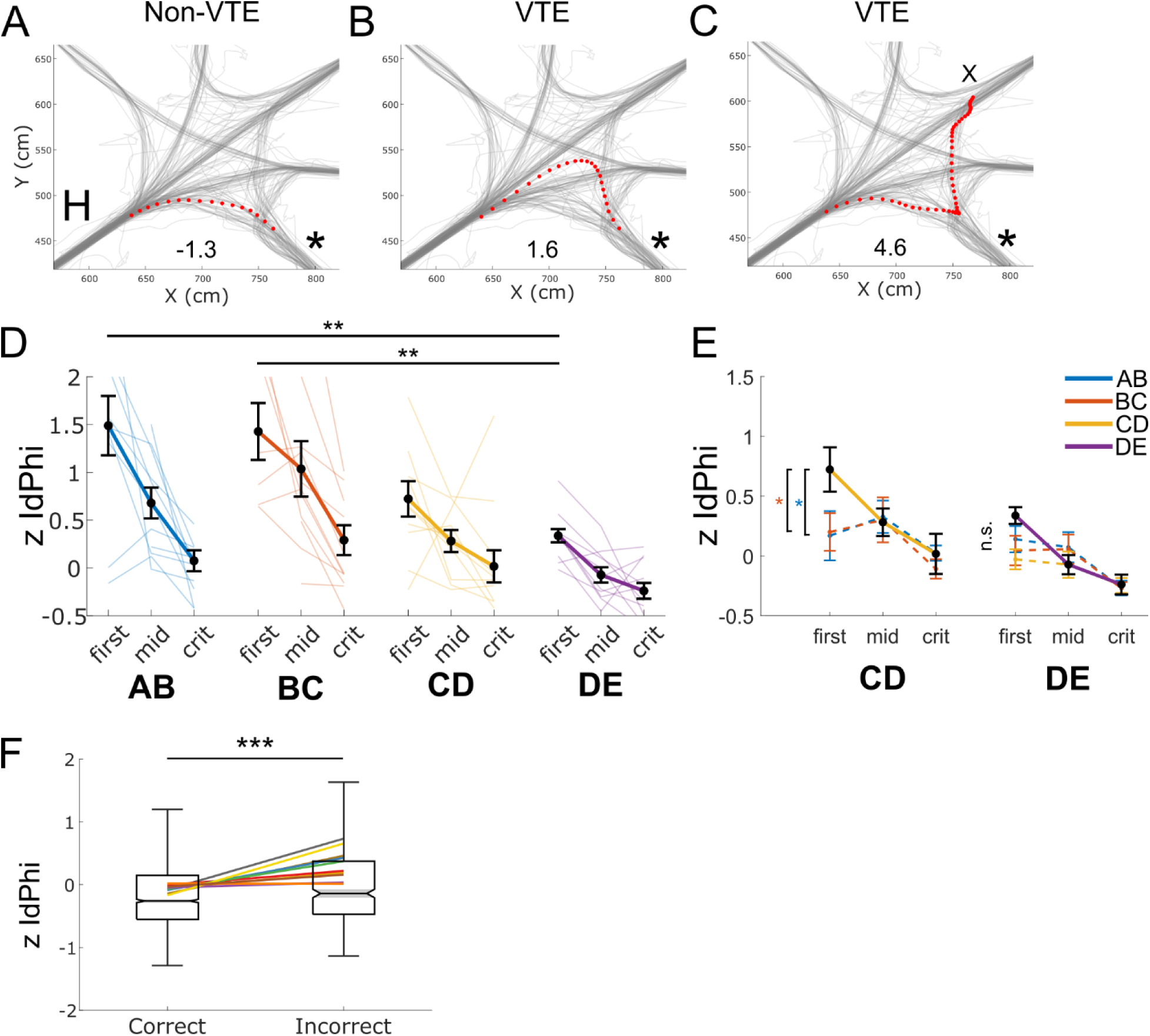
VTEs during premise pair training. **A-C)** Example choice trajectories from Home (H, bottom left) to the arm of choice. All examples are the same trial type where the correct arm is noted by the *. The inset value indicates zIdPhi that was used to quantify VTE behavior. **A)** Smooth ballistic trajectory to the correct arm. **B)** Example VTE event where rat begins on path to wrong arm but reorients to correct arm. **C)** Example VTE event where rat stops at the correct arm to reorient and choose the incorrect arm denoted by the X. **D)** Average z-scored zIdPhi’s across training for each trial type (n = 10). The first training day, the final criteria day for a training phase, and a midpoint day were chosen. Note the reduction in VTE over training of a given premise pair, and the increase in VTE at the introduction of a new premise pair. **E)** VTE quantification for all premise pairs shown during the designated days for C-D and D-E. The plots for C-D and D-E are the same as D, and zIdPhi for A-B and B-C is shown overlaid on those days. Note that when a new premise pair was introduced, all previously learned premise pairs were also presented in that session in a pseudo-random manner (e.g., A-B and B-C are presented when C-D is introduced as a new premise pair; and A-B, B-C, and C-D are presented when D-E is introduced as a new premise pair). **F)** zIdPhi values for correct vs. incorrect trials across all premise pairs. Each colored line corresponds to the average values for each rat (n = 10), shown for visualization (main statistical testing used all trials from all rats). Box plot shows median as central value. n.s., not significant. *, p < 0.05; **; p < 0.01; ***, p < 0.001.

### Test Performance Compared to Initial Premise Pair Learning

Next, we investigated the evidence to ascertain whether rats were using inferential reasoning to solve B-D. First, we examined the time course of B-D performance on the first 10 trials (**Figure 3B**). Using a four-trial window with a two-trial overlap, we found that rats on average perform significantly better than chance level of 50% on the first two blocks of trials (p’s < 0.0125), and significantly better than 75% on the last two blocks of trials (p’s < 0.0125), indicating rapid high performance on B-D. Next, if rats are indeed using inference based on their task schema to solve B-D and A-E, they should do better on these test pairs compared to the first time that they were presented with premise pairs C-D and D-E, which had not yet been assimilated into the task schema. To investigate this, we compared how well rats performed on the first day they encountered C-D and D-E during training and their performance on test day for B-D and A-E. We excluded A-B and B-C for a fair comparison since the first day of phase one (when A-B and B-C were introduced) was the first day the rats were on the maze and likely no memory schema had been formed yet. We found that rats performed significantly different on the first day a trial type was presented (repeated measures one-way ANOVA, F(3,18) = 17.4, p < 0.0001; **Figure 3C**), and in particular did significantly better on the test pairs (B-D & A-E) compared to the premise pairs (C-D & D-E) on the first day they are presented (all p’s < 0.05). Overall, rats’ high performance on the test pairs indicates utilization of the memory schema of the value hierarchy, as opposed to trial-and-error exploration when new arms are added during premise pair learning.

### Vicarious Trial-and-Error Behavior During Premise Pair Learning

VTEs provide a valuable behavioral readout of spatial deliberation (Bett et al., 2012; Miles et al., 2024; Redish, 2016; Stout et al., 2022). Previous studies indicate that VTEs remain low before rats understand the layout of the maze context (i.e., exploration with no schema,), increase in rate and peak as animals use trial-and-error to learn rules and relationships (i.e., with deliberation, the schema is forming and being tested), and diminish as the task becomes habitual (i.e., habit formation, exploitation with reliable schema being used; for review, see Redish, 2016). However, sufficiently complex tasks may show relatively elevated levels of VTEs even after task proficiency is reached to reflect an ongoing need for deliberation between options. Our task is unique among many existing paradigms since arms D and E are added during training, necessitating schema assimilation and accommodation. For example, during phase 1 of training rats could simply solve B-C by avoiding C. However, starting on phase 2, rats now need to update their representation of C to be context dependent (e.g., avoid C for B-C but choose C for C-D). We were therefore interested in the progression of VTEs over training and how adding new premise pairs altered VTEs.

We measured VTEs using the zIdPhi metric of integrated absolute value of the angular head velocity (Papale et al., 2012); for standardization, since all choice arms are in different locations at different angles from the Home arm, trajectory zIdPhi metrics were z-scored independently for each arm for each animal (see Methods). Any choice trajectory lasting longer than 2 seconds in the central choice zone was removed to avoid including trials where behaviors like grooming or excessive exploration occurred in the center zone. After removing these trials, the mean choice time was 0.75 seconds (± 0.24 S.D.). Three example trajectories from the same session and the same trial type are shown in **Figure 4A-C**. A ballistic, correct trajectory with no VTE can be seen in **Figure 4A** while **Figure 4B & 4C** show correct and incorrect VTE trials respectively. We measured the zIdPhi values across training, using the same session timepoints of “first”, “mid”, and “criterion” sessions as **Figure 2B**. As expected, zIdPhi values and hence VTE behaviors peak early on (A-B and B-C “first”) when uncertainty is high and gradually diminish with task proficiency. ZIdPhi values increase again when new arms / premise pairs are added and show similar decreasing trends during the course of learning new premise pairs.

We tested whether zIdPhi values were different on the first day of each premise pair introduction and found a significant difference in trial types for zIdPhi values on the first day of presentation (F (3,31) = 4.71, p = 0.008). Post-hoc multiple comparisons showed zIdPhi values were significantly higher on the first day of A-B and B-C compared to D-E (p = 0.02, p = 0.02, respectively). We also wanted to test whether VTEs increased for all trial types when a new arm was added. Thus, we focused on zIdPhi values for the first day of C-D and D-E (**Figure 4E**). On the first day of C-D, zIdPhi values were significantly higher for C-D trials than for A-B and B-C (F(2,27) = 5.1, p = 0.013). In contrast, on the first day of D-E, zIdPhi values were not different across the four trial types (F(3,36) = 1.8, p = 0.16). Taken together, the decrease in the zIdPhi of trials over training, notably peak values during introduction of new premise pairs, especially by D-E’s introduction, suggests that most rats have an established schema of the task by the time D-E is introduced that can readily assimilate new premise pairs, and novel D-E trials do not especially evoke additional VTEs due to uncertainty.

An increased presence of VTEs has been reported to be associated with both correct (Kidder et al., 2021; Miles et al., 2024) or incorrect trials (Hu & Amsel, 1995; Rosenblum et al., 2025; Santos-Pata & Verschure, 2018; Schmidt et al., 2013; Stout et al., 2022). The ambiguity may be due to learning time courses, how different tasks are performed, and their difficulty levels. We tested zIdPhi values between correct and incorrect trials via bootstrapping by down sampling with replacement the correct trials to match the number of incorrect trials. The first three days of training on phase one for all animals were not included, as rats were acclimatizing to the novel maze and learning the task structure, and had high error rates. In this transitive inference task, we found that incorrect trials had significantly higher zIdPhi values than correct trials (mean p value = 0.0002 (95% CI: < 0.0001, 0.0011); mean effect size = -0.24; 95% CI: -0.29, -0.20 **Figure 4F**). We also tested each rat individually to determine if their zIdPhi values were different across correct and incorrect trials. Eight out of the 10 rats had mean p-values less than 0.05. Thus, higher zIdPhi values are strongly associated with incorrect trials in this task.

### Vicarious Trial-and-Error During Inferential Reasoning

VTEs have long been regarded as a behavioral hallmark of deliberation, especially during challenging tasks that may rely on memory schema (Redish, 2016). We were interested in how VTE-like behavior occurred during inference testing when rats (N = 9, all tested rats) were challenged with inference trials. We examined the first five trials of B-D, as well as B-C, on test day when deliberation was likely highest (**Figure 1A-I**). We highlight B-C trials as they require the same trajectory to solve as B-D, but rely on retrieving a learned premise pair on test day rather than inferring the correct choice. We observed a diverse range of trajectories across rats and across trials for test pair B-D, shown for individual rats for initial 5 trials in **Figure 5**. A subset of animals (n=3, **Figure 5A-C**) made no errors and generally had ballistic, low zIdPhi trajectories. Another group of animals (n=4, **Figure 5D-G**) chose incorrectly on the initial trial or two, often with an elevated zIdPhi (**Figure 5E-G**), before getting the remaining trials correct with a lower zIdPhi. One animal (**Figure 5H**) showed high zIdPhi trajectories suggestive of high uncertainty, and high rate of error. The last animal shown in **Figure 5I** failed the inference test pair B-D, averaging 33% correct for the entire session, and appears to incorrectly choose D with ballistic trajectories on most trials.

**Figure 5:**
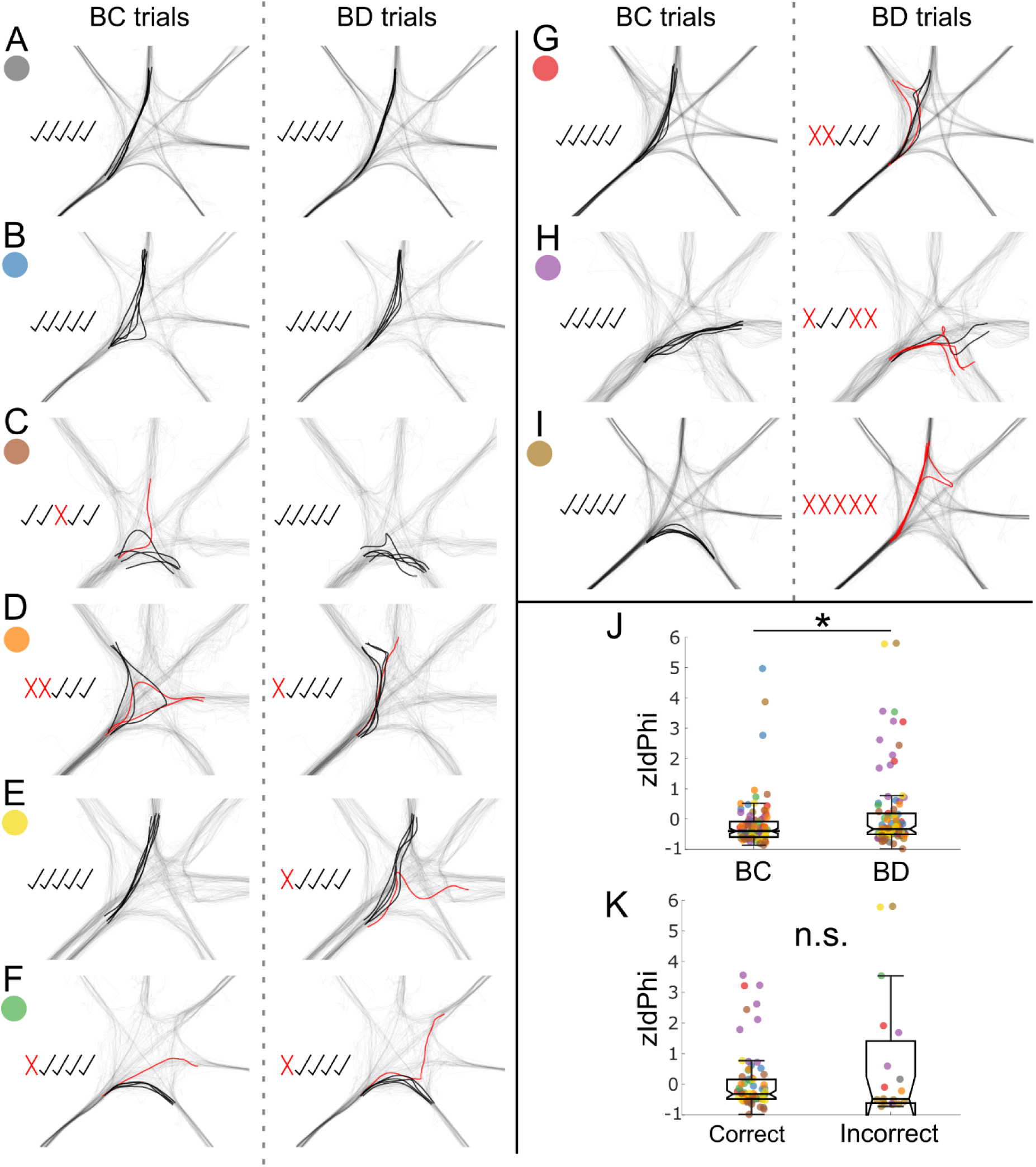
VTEs on inference test day. **A-I**) The trajectories for the first 5 trials of B-C (left columns) and B-D (right columns) for each rat (rows; colored dot corresponds to rats’ color across graphs). Incorrect trials are in red while correct are in black. Choice trajectories are overlaid on the tracking from the full session, shown in black with a low opacity. Inset check marks denote correct trials and X’s denote incorrect trials in chronological order for the corresponding trial type. Rats show a high variability in their choice trajectories with VTE-like-behavior not always being present during B-D inference choices. **J)** zIdPhi values for the first 10 B-C trials compared to the first 10 B-D trials for all tested rats on test day. Rats had a significantly higher zIdPhi value for B-D trials compared to B-C **K)** zIdPhi values for the first 10 B-D trials comparing correct and incorrect trials. Each color represents data from one rat, colors consistent across all figures. Box plots show median as central value. n.s., not significant. *, p < 0.05.

Restricting analyses to the first 10 trials of the two trial types on the test day, we found that zIdPhi values were significantly higher on B-D trials compared to B-C trials (linear mixed-effect model with rat ID and trial number as random factors; F(1,138) = 5.0, p = 0.03; **Figure 5J**). Next, we tested whether ZIdphi values were higher on correct trials compared to incorrect trials for B-D across the first 10 trials on test day, which may indicate that rats show deliberation-like behaviors (high ZIdphi) before making correct choices. We found no significant difference in zIdPhi values for B-D correct compared to incorrect trials for the group of rats that showed successful inference (n = 7; bootstrap Wilcoxon rank sum, p = 0.68, 95% CI: 0.23, 1.0; **Figure 5K**). However, the individual rat variability in performance and choice behaviors (zIdPhi) precludes an overall interpretation regarding deliberation during inferential reasoning.

The individual variability suggests that the inference strategy may be determined by the strength of schemas and retrieval ability from schemas for deduction. Overall, we find that VTE-like behavior is unique to individual animals across cognitive conditions (recall vs. inference) and performance (**Figure 5A-I**). We hypothesize that animals that can retrieve associations from existing schemas formed during learning with high accuracy will show less VTE behavior, whereas incorrect recall or delays in recall or will result in higher VTE behavior and deliberation during initial inference trials. Failure of inference tests likely indicates a lack of schema formation, with animals performing tasks based on learning of individual pair associations (A>B, B>C, etc.), without forming an overall schema structure. These hypotheses provide an avenue for physiological investigation, leveraging individual variability to assess links between potential physiological correlates of schemas and recall with behavioral performance.

## Discussion

We demonstrate the feasibility of a spatial transitive inference task for rats utilizing an automated radial maze. A majority of the animals trained on the task were able to learn the premise pairs and reach high levels of performance, and only two of the nine trained animals failed the inference test. We investigated deliberation-like behavior by analyzing vicarious trial and error events, quantified using zIdPhi values, during training and testing. We found a marked drop in zIdPhi values over training as rats became proficient. Adding a new premise pair early on resulted in high zIdPhi values for that pair, but adding a second new premise pair did not increase its zIdPhi values beyond the previous premise pairs. On test day we found zIdPhi values were highly variable across rats and performance. We found a modest increase in zIdPhi values for B-D trials over B-C. Overall, our data indicate that VTE’s are common during inference decisions and may indicate that utilization of a well-established schema allows animals to make novel choices without any deliberation.

The most common rodent TI paradigm utilizes odors as the stimuli mapped to the items of the value hierarchy, where odors are mixed into sand and placed in pots that rats must dig for a reward. While these studies were critical to establishing rodents’ abilities to perform transitive inference (Davis, 1992; Roberts & Phelps, 1994; Van Elzakker et al., 2003) and the brain regions necessary for inferential reasoning (DeVito, Kanter, et al., 2010; Dusek & Eichenbaum, 1997; Van der Jeugd et al., 2009), there are many drawbacks. Manual baiting and setup of every trial leads to lower throughput and could introduce variability to experimental conditions (e.g., depth of reward, pot positions). It is also ambiguous to determine the precise time period when rats may be perceiving the odors and making an inference judgment. Our version of the TI paradigm addresses these issues by using an automated maze and having each arm represent an item in the value hierarchy. It’s important to note that while our study uses space (the distinct arms) as the items, the spatial layout is not inherently related to the value hierarchy, as was the case in some rodent TI tasks (Roberts & Phelps, 1994). The radial maze also inherently has a decision area in the center of the maze where rats can perceive the open arms and make their decision. Furthermore, a rat’s choice of an arm is easier to quantify than the traditional time digging metric. Our new task, coupled with precise position tracking and electrophysiological recordings, will give valuable insights into the neuronal mechanisms of transitive inference such as the role of theta and replay sequences in schema formation and decision making. Such studies would also allow the comparison of rodent neural activity to non-human primates and humans during inferential reasoning to investigate shared and distinct mechanisms, such as common neural manifolds (Ramawat et al., 2023).

Despite the differences of our spatial TI task and training schedule from the traditional scented-pot version, the rats’ behavior and performance are remarkably similar to the existing literature. Across studies, premise pair acquisition failure rates are similar, with about 20-25% of the rats trained failing to learn all premise pairs on five-item sets (Davis, 1992; Dusek & Eichenbaum, 1997; Silverman et al., 2015; Van Elzakker et al., 2003). In our task, rats generally end on higher performance levels (∼90%) compared to the odor tasks (∼80%) despite the performance criteria being the same for both tasks (∼75%). Surprisingly, rats in this spatial task do not appear to have the typical “V” shaped performance curve. Generally, rodent and NHP (Brunamonti et al., 2016) subjects’ average performance is highest for the end premise pairs (e.g., A-B, D-E) and lowest for the “inner pairs” (e.g., B-C, C-D). This is likely due to the end pairs having a simple rule to get correct, i.e., always pick A and never pick E. The lack of V-shaped performance on our task could be attributed to a reward provided on arm E for correct D-E trials. In contrast, in traditional paradigms, a reward is never encountered on the end item. Thus, rats in our task may have a higher propensity to choose arm E on D-E trials as rewards have been encountered there.

In addition, it appears that training durations are remarkably similar across rodent TI paradigms. In our paradigm of intermixed premise pairs, rats took an average of 21 days to complete training. Intermixing trials is more common in NHP (Gillan, 1981) studies compared to rodent ones, with previous reports that rats cannot learn with intermixed training (Donald, 1987). Previous studies using blocks of trials that transition into intermixed report training time in days were similar at 26 (DeVito, Kanter, et al., 2010) and 18 (DeVito, Lykken, et al., 2010). Interestingly, a touchscreen version of the task also took 22 days of training (Silverman et al., 2015). However, our task required an order of magnitude more trials, averaging 2,215 (737 S.D.), compared to block design tasks. The training regime of most other TI tasks is the presentation of blocks (usually starting around 10 trials long) of each premise pair until the criterion is reached and the block size is reduced until the final phase of intermixing. Where reported, rats required 300-400 total trials to learn the odor-based version of the task (Dusek & Eichenbaum, 1997; Van der Jeugd et al., 2009). Based on the reported training methods, we estimate the studies reporting training days had similar trial counts of a few hundred (DeVito, Kanter, et al., 2010; DeVito, Lykken, et al., 2010). Thus, while total training days may be similar, it’s likely more efficient in terms of total trials to train rats with blocks of specific premise pairs before transitioning to intermixed presentation. Furthermore, these differences raise important questions on the most efficient way to build memory schemas.

To our knowledge, there is only one other rodent spatial transitive inference task that was developed for mice (Abdou et al., 2024). In this version of the task, a corridor has five distinct (color, geometry, and texture) “rooms” flanking each side. Each context was assigned an item on the hierarchy (A>B>C>D>E) and mice made their choice between two rooms by staying in one of them for 10 seconds. However, direct comparisons to this paradigm are challenging; the training schedule in their task consisted of only two premise pairs a day with no intermixing of all four premise pairs in one training session. Furthermore, mice in this task failed the first inference test and only successfully inferred after sleeping and being retested. As a result, we hesitate to make direct comparisons to their valuable study.

Unlike previous rodent TI paradigms, our radial maze version has a clear deliberation area that can be used to investigate potential reasoning behaviors such as vicarious trial and error events. In addition, by serially adding premise pairs, we have distinct time points where the task gets more challenging. Traditionally, VTEs have been ascribed to reflect the indecision of deliberation as rodents vicariously sample potential choices (Redish, 2016). Furthermore, it has been demonstrated that VTE rates peak during learning and diminish as choices become automated. Neural recordings in spatial decision-making tasks have demonstrated the relevance of the hippocampal-prefrontal circuit in deliberative behaviors and VTEs (Amemiya & Redish, 2018; Bett et al., 2012, 2015; Hasz & Redish, 2020, 2020; Johnson & Redish, 2007; Kidder et al., 2021; Miles et al., 2024; Papale et al., 2016; Rosenblum et al., 2025; Schmidt et al., 2019; Stout et al., 2022). However, recent studies have posited that VTE behaviors are heterogeneous and may not always reflect deliberation (Miles et al., 2024). Miles reported high VTE during periods of learning that coincided with correct trials, aligning with the notion of VTE’s as a readout of deliberation. A separate time of high VTEs was found when rats were not learning and getting trials wrong, possibly reflecting a behavioral state akin to uncertainty. We find that high zIdPhi values, a quantification of VTEs, are highest early in training and diminish as rats become proficient. Adding in the first additional premise pair, C-D, results in high zIdPhi values. However, this elevation is not seen when D-E is added, potentially showing that adding E evokes less uncertainty than adding D. The lower zIdPhi values for D-E may also reflect a more efficient schema assimilation process for D-E, as the rats have more experience with the task compared to when C-D is added. On test day, we find that, on average, zIdPhi values are higher for B-D trials compared to B-C, despite rats doing equally well on both. Thus, rats may have higher zIdPhi values while inferentially reasoning on B-D trials and having lower zIdPhi values for the well-trained B-C trials. However, many rats had low zIdPhi’s value and successfully inferred. This may indicate that some animals with well-established schemas can make inferential decisions rapidly with no deliberation necessary. Overall, these findings support the notion that VTEs occur during periods of learning but are not a reliable behavioral correlate of inferential reasoning.

The immediate utility of VTEs in decision making is also unclear. If VTEs primarily reflect deliberation, one might expect that after a VTE, a rodent will have a higher probability of making a correct choice (Kidder et al., 2021; Miles et al., 2024). However, we find that zIdPhi values are consistently higher on incorrect trials than on correct trials both during training and testing, similar to other studies (Hu & Amsel, 1995; Rosenblum et al., 2025; Santos-Pata & Verschure, 2018; Schmidt et al., 2013; Stout et al., 2022). This may further indicate that during schema-based tasks, VTEs can be associated with periods of indecision when rats are more likely to make errors. Overall, we further bolster the notion that VTEs can occur under different experimental conditions and cognitive states and further investigations into their cause and purpose are needed, including neurophysiological investigation during behavioral periods of uncertainty and deliberation, which may reveal differences.

To conclude, we have demonstrated the feasibility of a spatial transitive inference task for rats that results in similar inference test success compared to the common odor-based TI paradigm. Our version of the task, with sequential training and deliberation zone, is more amenable to physiological recordings which are needed to investigate the neural mechanisms underlying inferential reasoning and memory schema. Computational model investigations into the potential mechanisms of solving transitive inference have demonstrated that numerous artificial neural networks can successfully solve TI tasks (Kay et al., 2024; Miconi & Kay, 2025), including hippocampal models (Frank et al., 2003; Kumaran & McClelland, 2012; Wu & B Levy, 2001; Wu & Levy, 1998). We expect that this spatial version of the classic transitive inference task, coupled with physiological recordings from the hippocampus and prefrontal cortex, can aid in elucidating the mechanisms supporting memory schema and inferential reasoning.

