## Supplemental Figure 1 for "A Novel Rat Spatial Inference Task Reveals Rapid Deduction of Transitive Relationships Using Schemas and Deliberation"

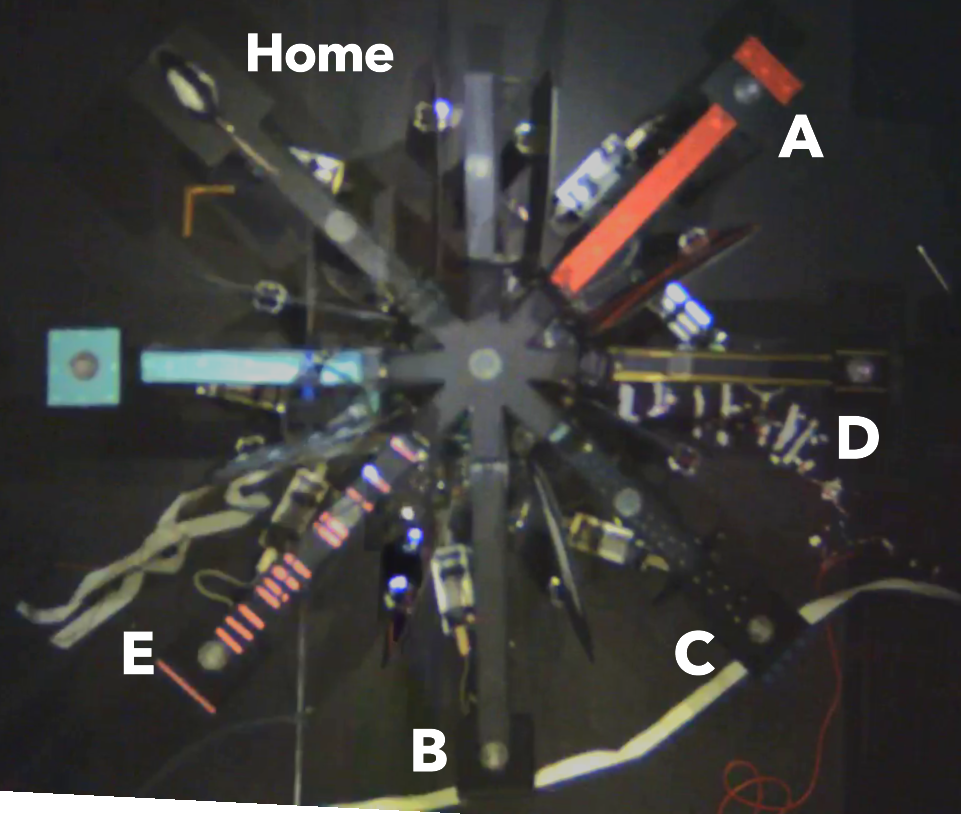
**A)**

|  | **Arm - Item assignments** | | | | | | | | | |
| --- | --- | --- | --- | --- | --- | --- | --- | --- | --- | --- |
| **Arm Number** | **BP13** | **BP15** | **BP16** | **BP19** | **BP21** | **BP22** | **TH405** | **TH508** | **TH510** | **TH605** |
| **1** | Home | Home | Home | Home | Home | Home | Home | Home | Home | Home |
| **2** | ~~-~~ | ~~-~~ | ~~-~~ | ~~-~~ | ~~-~~ | ~~-~~ | ~~-~~ | ~~-~~ | ~~-~~ | ~~-~~ |
| **3** | C | B | A | A | C | B | D | A | A | B |
| **4** | B | E | D | C | D | C | B | C | D | E |
| **5** | D | C | C | E | E | D | E | E | C | A |
| **6** | A | D | B | B | B | E | A | D | B | C |
| **7** | E | A | E | D | A | A | C | B | E | D |
| **8** | ~~-~~ | ~~-~~ | ~~-~~ | ~~-~~ | ~~-~~ | ~~-~~ | ~~-~~ | ~~-~~ | ~~-~~ | ~~-~~ |

**B)**

**Supplemental Figure 1: Maze depiction and item-arm assignments. A)** Overhead view of the maze with arm designations for one rat (BP16 in **Supp Figure 1B**). Unique contexts can be seen for each arm, such as red Polyvinyl Chloride shelf liner (arm A), golden vinyl stripes (arm D), hot glue dots (arm C), black foam (arm B), and pink horizontal stripes (arm E). Complimentary cues were also on the walls between arms. The arms 45° from Home, i.e. arm #2 and #8, were not used. **B)** Arm-item assignments for all rats used in the study. Arm-item assignments were randomly generated (using the randomization tool in Google Sheets) prior to premise pair training. Any configuration with a simple egocentric strategy to solve was rejected and regenerated. Arm-item configurations stayed constant for the whole experiment.
